# Viscoelastic RNA entanglement and advective flow underlie nucleolar form and function

**DOI:** 10.1101/2021.12.31.474660

**Authors:** Joshua A. Riback, Jorine M. Eeftens, Daniel S. W. Lee, Sofia A. Quinodoz, Lien Beckers, Lindsay A. Becker, Clifford P. Brangwynne

## Abstract

The nucleolus facilitates transcription, processing, and assembly of ribosomal RNA (rRNA), the most abundant RNA in cells. Nucleolar function is facilitated by its multiphase liquid properties, but nucleolar fluidity and its connection to ribosome biogenesis remain unclear. Here, we used quantitative imaging, mathematical modeling, and pulse-chase nucleotide labelling to map nucleolar rRNA dynamics. Inconsistent with a purely diffusive process, rRNA steadily expands away from the transcriptional sites, moving in a slow (~1Å/s), radially-directed fashion. This motion reflects the viscoelastic properties of a highly concentrated gel of entangled rRNA, whose constant polymerization drives steady outward flow. We propose a new viscoelastic rRNA release model, where nucleolar rRNA cleavage and processing reduce entanglement, fluidizing the nucleolar periphery to facilitate release of mature pre-ribosomal particles.

---

Cells compartmentalize biomolecules into organelles to enable spatiotemporal control over the formation, processing, and regulation of macromolecular complexes essential for life. An important class of such compartments is biomolecular condensates, which despite their lack of a bounding membrane are coherent and often liquid-like, with roughly spherical shapes, some degree of internal mixing, and exchange with the surroundings (*1–3*). These dynamic features are facilitated through weak multivalent interactions, typically involving oligomerized intrinsically disordered regions (IDRs), that drive liquid-liquid phase separation (LLPS), and provide an internal cohesivity to the constantly restructuring biomolecular network (*4–7*).

Despite a plethora of studies supporting the LLPS concept, the exact physical nature of biomolecular condensates has been debated (*8, 9*). This sometimes arises from over-simplified representations, including the binary characterization of condensates as either purely liquid-like, or else reflecting some qualitatively different assembly process. Such simplifications ignore the inherent biological complexities of intracellular condensates, including their polymeric nature (*10*), compositional complexity (*11*), and the role of active processes (*12*). For example, condensates typically contain hundreds of different components, which contribute to the thermodynamic driving forces of phase separation in ways that are not captured with two-component mean-field models (*13–15*). A full picture of how condensates contribute to biological function will require an understanding of how they differ from simple liquids, and how those properties are exploited by cells to impart biological function.

One of the most fascinating and still poorly-understood cellular processes occurs in the nucleolus, the most prominent nuclear condensate, which is the site of ribosome subunit assembly in all eukaryotes (*16*). The nucleolus consists of three distinct and concentrically-ordered layers, the FC (fibrillar center), DFC (dense fibrillar component), and GC (granular component), which roughly correspond to the sites of ribosomal RNA (rRNA) transcription, processing, and assembly into the small (pre-40S) and large (pre-60S) ribosome subunits, respectively (*17*). These distinct layers are thought to represent immiscible sub-phases, which each form via LLPS, but do not mix with one another and remain nested, due to their differential surface tensions (*18–21*). These immiscible phases have been suggested to facilitate the sequential processing of rRNA transcripts, possibly through a “hand-off” mechanism, whereby sequential processing steps occur in an assembly-line (*22*). However, it is unclear how this can occur within a liquid environment, and how the material properties (i.e. viscosity, surface tension) of the different immiscible layers might contribute to the intricate multistep process of ribosome biogenesis.

To gain better insight into how the material properties of the nucleolus facilitate its function, we first re-examined some of the classic dynamic features of nucleoli, which have been used to argue for its simple (i.e. Newtonian) liquid properties. Simple liquids and solids are differentiated by indicators of their mesoscale dynamics, with liquids showing rapid diffusion, coalescence, and rounding to minimize surface area **(Figure 1A)**. Consistent with previous studies (*19, 20, 23, 24*), the nucleolus exhibits fast (<1min) internal mixing of the scaffold protein NPM1 **(Figure 1B, D)**, but slow fusion and frequently aspherical shape in cells **(Figure 1E,F,H)**, suggestive of both liquid-like and solid-like features; interestingly, continuously dividing cells (e.g. U2OS, iPSCs) tend to have less spherical nucleoli than post-mitotic rat neurons cells (**Figure 1H**). In contrast, engineered phase separation of multivalent disordered proteins (*7*), optogenetically induced within the nucleus, results in spherical condensates with fast internal mixing and fusion **(Figure 1C,D,E,G)**, both indicative of a classical liquid. Thus, neither the intracellular milieu (e.g. chromatin), nor the proteinaceous scaffolding of biomolecular condensates is sufficient to explain the unusual material properties of nucleoli.

**Figure 1.**
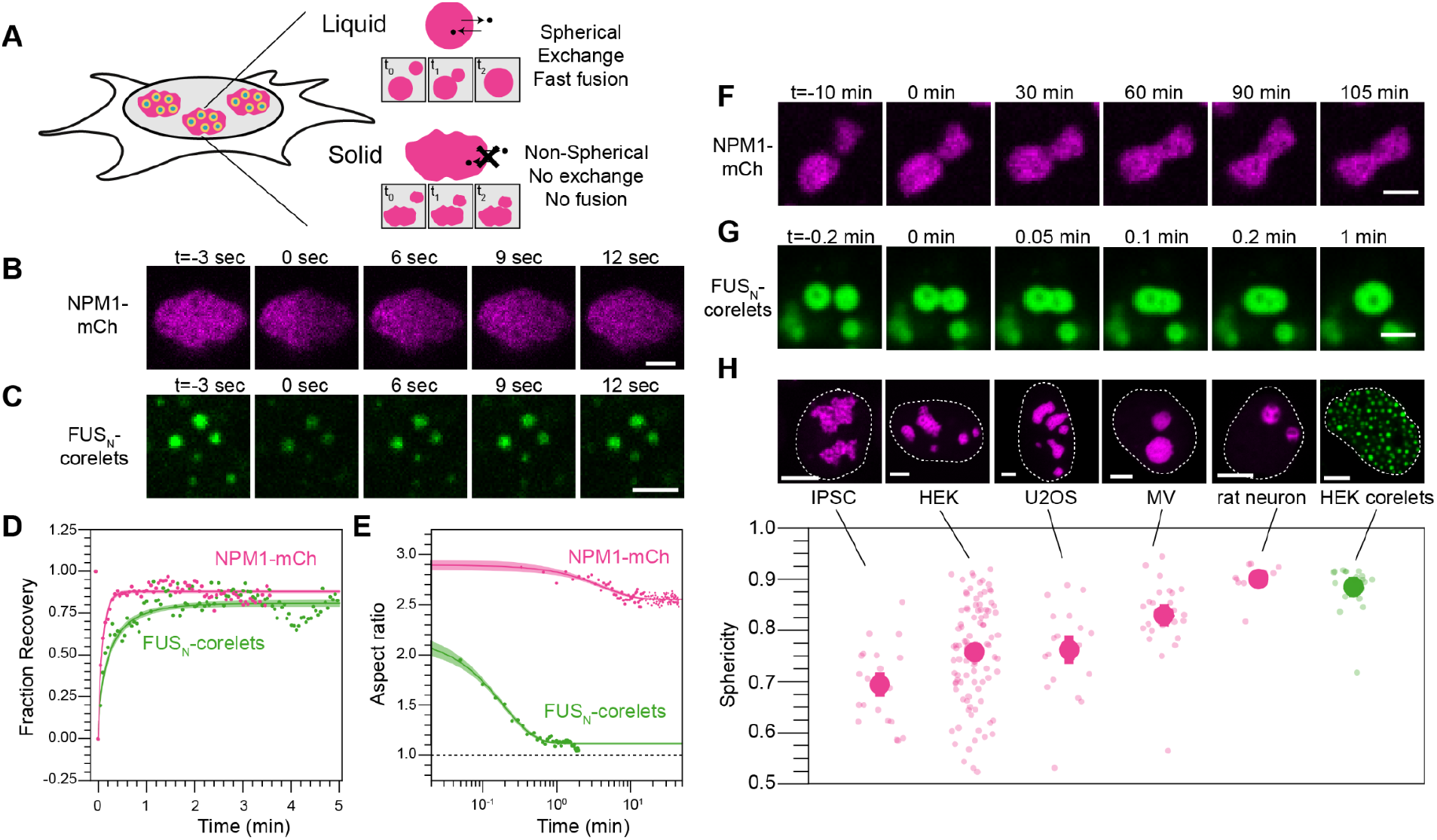
The nucleolus exhibits complex material properties, while synthetic condensates behave like typical liquids. All data are in HEK cells and all scale bars are 2µm unless otherwise noted. A) Schematic detailing expected properties of liquid-like vs solid-like nucleoli. B) Half FRAP of NPM1-mCh in the GC demonstrates rapid recovery. C) Similarly, FRAP of multiple small engineered droplets (FUS_N_-mCh-sspB co-expressing with NLS-GFP-FTH-iLiD, (*7*)) recovers quickly after photobleaching. D). Quantification and averaging of FRAP traces for both NPM1 (half FRAP of individual nucleoli in N=4 nucleoli) and FUS_N_ (whole FRAP followed by normalization for total fluorophore bleach in 90 condensates over N=8 cells) reveals recovery on a timescale of seconds. E) The aspect ratio of merging condensates shown in F and G demonstrates rapid kinetics for engineered droplets, but no rounding on a timescale of hours for nucleoli. F) Nucleoli, marked by NPM1-mCh, fuse but fail to fully round even on long timescales. G) Engineered droplets fuse and round rapidly upon contact. G) The average morphology of nucleoli ranked from lowest to highest sphericity and engineered droplets. Scale bars are 5µm. H) Morphology is quantified by sphericity, given by 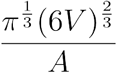 where V and A are the volume and area, respectively; deviation from 1 indicates nonspherical shape. Error bars show cell-average standard error of the mean with smaller transparent points showing the individual volume-averaged nucleoli for each cell analyzed, or each droplet analyzed for the case of HEK corelet. The number of cells (nucleoli) analyzed are 23 (68), 99 (803), 20 (81), 28 (42), and 13 (38) for IPSC, HEK, U2OS, MV, and rat neurons, respectively.

These observations suggest that the nucleolus may not reflect a simple fluid, and instead could exhibit some degree of viscoelasticity. Viscoelasticity is common in soft condensed matter physics, where so-called complex fluids exhibit material properties of both liquids and solids (*25*). Complex fluids include polymeric liquids, where long serpentine polymer chains are interwoven with one another. This polymer entanglement leads to elasticity on short timescales, while on long time scales polymer chains slide past one another, causing viscous relaxation (*25*). Associative polymers that stick to one another can give rise to additional elasticity or even gelation, resulting in time-independent, solid-like behavior (*26, 27*). In the context of biomolecules, several recent studies underscore how RNA-RNA and RNA-protein interactions can result in solid-like material properties (*28–31*). rRNA is produced within the nucleolus at high concentrations (*32*), where each rRNA is several kb long, and interacts with various other nucleolar biomolecules; remarkably, the 13.3 kb-long nascent chain corresponds to a total extended contour length of roughly 10 µm, several times larger than the typical diameter of the entire nucleolus (**Figure 2A**)(*33*). We reasoned that such long and highly concentrated polymeric chains would likely entangle with one another (**Figure 2A**). Consistent with this concept, a rough estimate suggests that rRNA concentrations are 10-fold greater than the overlap concentration (**Figure 2A, Supplemental text**), at which separate polymer chains begin to entangle with one another (*27*).

**Figure 2.**
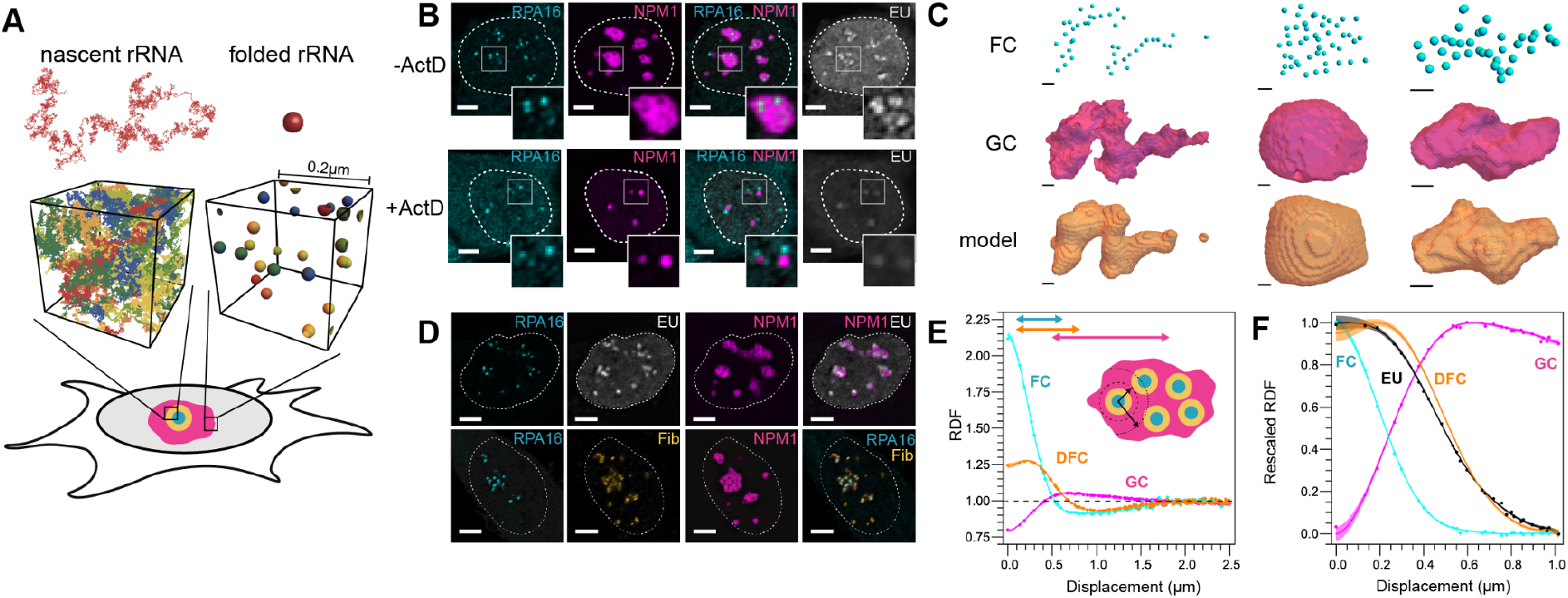
Entangled rRNA dictates nucleolar shape. A) Modeled as a random coil, nascent rRNA is ~15 fold larger in diameter than a folded rRNA subunit, resulting in substantially different degree of overlap between different chains within a calculated volume of the nucleolus, using rRNA transcript density estimated from experimental measurements (**Supplemental Text**). B) Inhibition of RNA transcription by ActD results in morphological changes of the nucleolus. Scale bar is 5µm. C) Within a simple shape model (**Supplemental Text**), FC location is sufficient to describe nucleolar shape in HEK, MV, and U2OS (left, middle, right). Scale bar is 1µm. D) RNA (EU; white) is transcribed at the interface of FC (RPA16; cyan) and DFC (Fibrillarin; yellow) (Scale bar is 5µm. E) The RDF of nucleolar phases follows concentrically (N=131 nucleoli). F) rRNA is primarily located in the DFC 30 minutes after addition of EU to the media (N=198 nucleoli). Note the DFC RDF fit is from D.

To examine the concept that entangled rRNA influences nucleolar shape, we partially inhibited Pol1 transcription by the addition of the drug Actinomycin D. Consistent with previous studies (*34, 35*), nucleoli become more round, and develop “caps” **(Figure 2B)**. The role of rRNA transcripts in controlling nucleolar shape is further supported by our finding that, within a simple transport-inspired model, the location of the FC (determined by monitoring RPA16, a subunit of Pol1I) is sufficient (R^2^ = 0.96 ± 0.01) to describe the overall shape and asphericity of nucleoli, prior to inhibition (**Figure 2C,S1A-B**). Taken together, these data support the concept that the constant transcription and formation of a viscoelastically-entangled rRNA gel strongly impacts nucleolar shape.

To obtain a high resolution, quantitative view of the relationship between rRNA and nucleolar structure, we examined nucleolar organization by calculating the radial distribution function (RDF), which is analogous to the pair-correlation function, *g*(*r*), commonly used to quantify the structure and dynamics of non-living materials (*36, 37*) **(Figure 2D,E,F)**. The RDF represents the relative intensity, radially averaged over a specified distance from a particular location, here the center of any FC within the nucleolus; an RDF greater than or less than unity is equivalent to a local spatial enrichment or depletion, respectively. As expected, the concentric nature of the FC (RPA16), DFC (Fibrillarin), and GC (NPM1) layers are clearly identified through calculation of the RDF **(Figure 2D,E)**. A local minima in the FC and DFC RDFs becomes apparent at ~1µm indicating half the average distance between the center of two FCs (**Figure 2E, S2A-D**).

We next sought to use this approach to quantify rRNA dynamics in the nucleolus. Observing native RNA dynamics in cells is challenging due to the lack of in situ RNA labeling. Early biochemical methods took advantage of the abundance of rRNA compared with other RNA transcripts, and followed nucleolar RNA via radioactive sulfur in uridine or BrdU (*34, 38*). Building from these early studies, we monitored rRNA through incorporation of nucleotide 5-ethynyl uridine (EU), which can be visualized following fixation and fluorescent modification of the EU (*39*). Following EU labeling for 30 minutes, images reveal EU signal localized at the DFC phase of the nucleolus. This is also captured by the calculated RDF for the EU signal, which closely coincides with the DFC curve (**Figure 2F**), consistent with previous results indicating RNA initially localizes at the FC/DFC interface (*21*). Thus, the RDF is a quantitative method to determine rRNA localization in the nucleolus.

We next sought to quantify the dynamics of rRNA transport, through a pulse-chase labelling protocol (**Figure 3A**). Upon incubating cells with EU for 30 min (“pulse”) followed by washout (“chase”) for varying times prior to fixation, we find that the RNA moved progressively away from the FC/DFC (**Figure 3B, S3A-D**). We quantified this data with the RDF analysis described above. The time-independent RPA16 and NPM1 RDFs indicated both a high degree of reproducibility of and a minimal impact of EU on nucleolar form during the experiment (**Figure S4A-B**). In contrast, as expected, the EU RDF changes with time, including a decrease and broadening of the peak of the RDF with time (**Figure 3C, S4D)**. Unexpectedly, however, the peak in the signal persisted, and moved radially at a slow apparent velocity of ~1Å/s with time, a clear indication of motion that is directed, rather than purely diffusional **(Figure 3C-E)**; indeed, solutions to the diffusion equation only exhibit a persistent and shifting peak when advection is included **(Figure 3C, right)**, while a simple partitioning model for movement between nucleolar subphases also fails to capture this motion (**Figure S4E-F**). These observations suggest that advection or flow, resulting from the large-scale polymerization of nucleotides into rRNA at the FC/DFC interface, drives slow directional motion of entangled RNA away from the FC/DFC.

**Figure 3.**
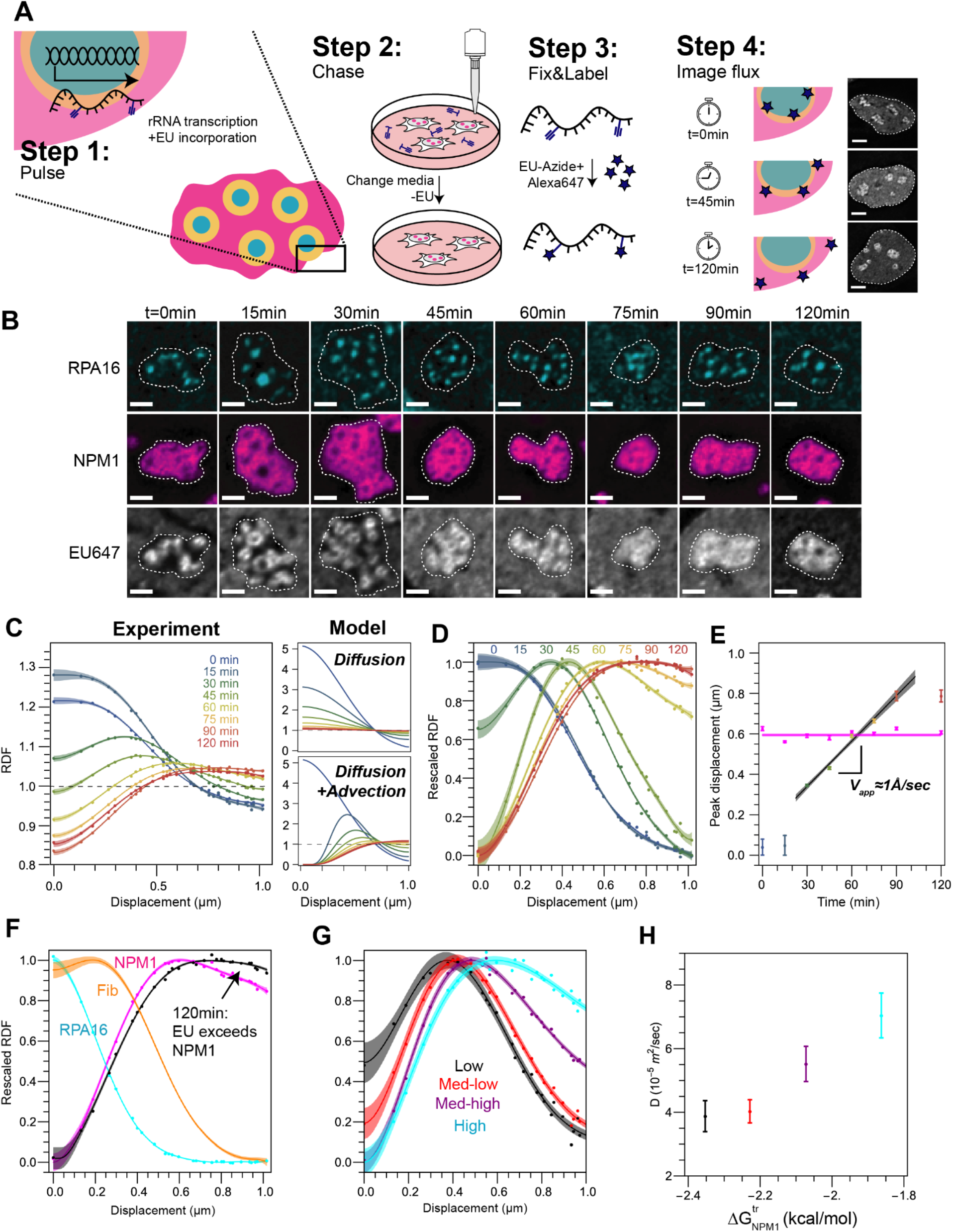
rRNA movement through the nucleolus reflects advective flow. A) Schematic of pulse-chase EU labelling experiment. Scale bar is 5µm. B) RNA (EU; white) moves away from its source (RPA16; cyan) progressively. Dashed lines demarcate individual nucleoli. Scale bar is 2µm. C) Left, quantification of RNA peak movement over time (N= 198, 155, 166, 159, 182, 279, 196, 201 nucleoli for sequential timepoints). Model, an analytical solution to the spherical advection-diffusion equation, of how the RDF would look in case RNA movement was driven by diffusion (top) or diffusion plus advection (bottom). D) Rescaled quantification of C. E) Maximum enrichment of RNA peak over time (timepoint coloring as in C, black line is fit). Maximum NPM1 enrichment remains stable over time (pink points and fit). F) Nucleolar phases and EU rescaled RDF at 120 minutes highlighting the EU peak exceeding the NPM1 peak in their respective RDFs. G) RDF at 45 minutes as a function of NPM1 concentration (N=120 nucleoli). I) Diffusion coefficients as a function of NPM1 concentration.

When characterizing transport phenomena, the relative strengths of advection compared to diffusion is quantified by the Peclet (Pe) number. In the limit of low Pe (Pe<<1), where diffusion dominates, the EU RDF should eventually reflect diffusive mixing throughout the nucleolus, or at least throughout the GC, if transcripts have a thermodynamic preference for partitioning into the GC (*13*). However, at the longest time probed, 120 minutes after labeling, the location of the peak of the EU signal extends ~0.2 um beyond that of the GC/NPM1 RDF (**Figure 3F**), suggesting a slight bias towards the GC periphery, detectable in the average EU signal. Moreover, by solving for a 3D-approximation to the advection-diffusion equation (**Figure S5J**), we determine a Pe number ~0.2 ± 0.1. These data indicate that the motion of rRNA reflects not only diffusive motion and thermodynamic partitioning, but also significant advective flow.

Taken together, our data suggests that rRNA forms a viscoelastic gel, whose constant polymerization at the FC pushes itself radially outward. Interestingly, this physical picture implies that as rRNA progressively matures and folds into compact ribosomal subunits, not only does its valence decrease (*13, 40*), but it should also become less entangled, and thus able to move in a more diffusive fashion; indeed a simple calculation suggests that fully processed and assembled rRNA chains will overlap with themselves over 3,000-fold less, compared with the nascent chains (**Figure 2A**; **Supplemental Text**). To test this concept, we explored whether the dynamics of rRNA transport are impacted by the relative concentration of NPM1, which binds rRNA and has been proposed to chaperone its folding and maturation (*19, 41*). We find that at higher concentrations of NPM1 (quantified as more negative ΔG^tr^ values, **Figure S5A**, methods(*13*)), the EU RDF peak shifted outwards, indicating substantially faster rRNA transport (**Figure 3G, S5**), even above the increase in the size of nucleoli. Fitting for the transport coefficients revealed a substantial, concomitant increase in diffusivity, concomitant with a slight decrease in the Peclet number, consistent with diffusivity becoming more dominant compared with advection (**Figure 3H, S5**); note the very low values of the fitted diffusion coefficient. Thus while key nucleolar proteins like NPM1 are much more highly dynamic than rRNA, their relative concentration appears to significantly impact rRNA entanglement and viscoelasticity, and thus rRNA transport through the nucleolus.

We next sought to directly test the concept of nucleolar viscoelasticity. The material properties of a viscoelastic microenvironment are typically quantified using passive microrheology, i.e., by measuring the mean squared displacement (MSD) of tracer probes, typically beads (*42*). However, such measurements within living cells are extremely challenging, for various reasons (*43*), including bulk motion of the cell/nucleus. Nonetheless, we sought to establish a microrheology-type measurement, by quantifying the pairwise MSD of FCs labeled by RPA16-GFP, i.e, < *d*^2^ >= < (*x*_*i*_ − *x*_*j*_)^2^ >_*t,i*≠*j*_for all pairs (*i, j*) of FCs in a particular nucleolus (*44*). We find that the pair MSD of nucleolar FCs is small and strongly subdiffusive on short timescales, even above the noise floor (**Figure S6**), and can be fit to a nonlinear Maxwell fluid model describing motion inside viscoelastic polymer melts (*45, 46*); by contrast, the linear pair MSD of optogenetically induced droplets in the nucleoplasm fit this model poorly (**Figure 4A**). On long timescales, the relative motion of FCs is more diffusive, with a diffusion coefficient of approximately 3 × 10^−5^ *μm*^2^/*s*. Interestingly, this number is remarkably similar to the calculated diffusion coefficient associated with the fluctuating component of the motion of nascent rRNAs (5 × 10^−5^ *μm*^2^/*s*) (**Figure 3H**). Assuming that the hydrodynamic radius of the FC is roughly comparable to that of nascent rRNA, this is consistent with the hypothesis that the nonlinear motion of the NORs is due to nucleolar viscoelasticity, arising from rRNA entanglement.

**Figure 4:**
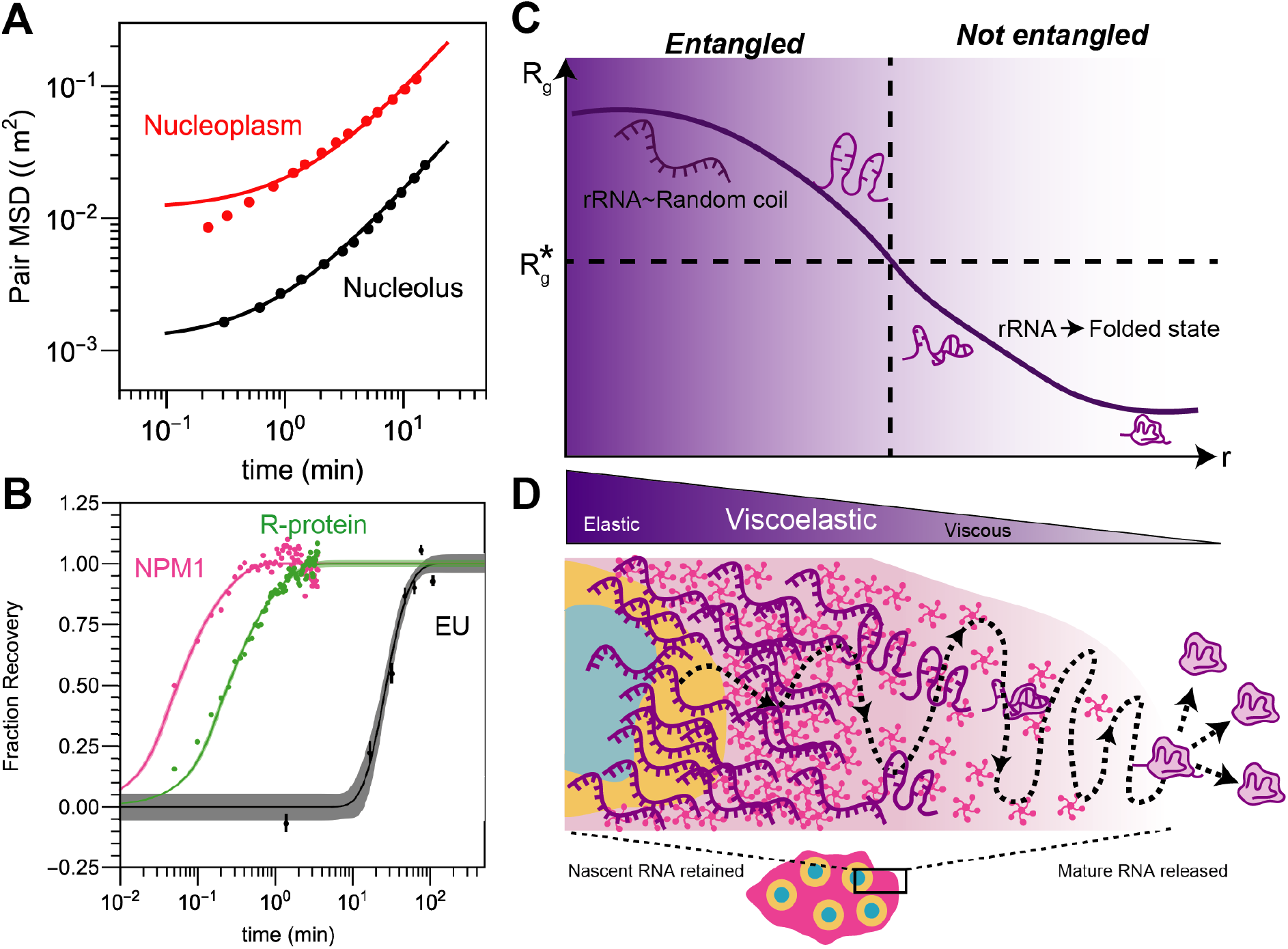
Nucleolar viscoelasticity and release of mature rRNAs. A) Pairwise MSD of long-time dynamics of FCs in HEK cells (‘Nucleolus’; 508 pairs of FCs in 7 nucleoli) and engineered droplets in U2OS nuclei (‘Nucleoplasm’; 31582 pairs in 18 cells, re-analyzed data from (*47*)). FCs fit a nonlinear Maxwell model, while engineered droplets do not. B) NPM1, R-proteins, and RNA have different dynamics. C) Schematic depicting changes in rRNA conformation, which leads to a progress decrease in the effective size of rRNA (i.e. Radius of gyration, R_g_), dropping below the size (R_g_*) at which chains are no longer overlapping and entangled. D) Schematic of nucleolar viscoelasticity during the progression of ribosome subunit assembly; mature pre-ribosomal particles move more diffusively through the peripheral GC liquid and are released into the nucleoplasm.

With nascent rRNA chains giving rise to strong entanglement and viscoelasticity, it may be unclear how nucleolar proteins such as NPM1 can exhibit rapid dynamics (e.g. FRAP recovery), which is often mistakenly taken as sufficient evidence of a liquid-like material state. Indeed, NPM1 exhibits rapid FRAP recovery with diffusion coefficient of 2 × 10^−1^ *μm*^2^/*s* orders of magnitude faster than that determined from our EU data (**Figure 4B**). Indeed, utilizing our EU data to calculate an equivalent FRAP recovery curve for rRNA (Methods), we find that the characteristic timescale for molecular dynamics is ~3 orders of magnitude slower for rRNA compared to NPM1; this is consistent with the vast differences in dynamics recently observed for nascent rRNA and NPM1 mixtures *in vitro* (*31*). Thus, while rRNA forms a slowly-relaxing viscoelastic gel that dominates the bulk material properties of the nucleolar interior, nucleolar proteins are only interacting with this gel transiently, and can apparently diffuse readily through its interstices (*48*).

Importantly, in deriving the apparent rRNA FRAP curve, we are effectively averaging over all processing/conformational states, and are likely dominated by the slow, entangled nascent chains, with the mobility of mature, folded chains being significantly faster. Consistent with this concept, the ribosomal protein RPL5, which despite being smaller exhibits a slightly lower mobility compared to NPM1 (**Figure 4B)**, suggesting incorporation into the RSU (*49*) is nonetheless several orders of magnitude more mobile compared with the average RNA signal. Thus, as rRNAs progress farther into the GC and fold together with ribosomal proteins, they become significantly smaller, and less kinetically trapped through entanglements, giving rise to a more fluid-like material state (**Figure 4C)**. These more mature ribosomal subunits also exhibit decreased interaction valence with other nucleolar components, and are thus ultimately subject to thermodynamic expulsion from the nucleolus (*13*).

Our findings provide a dramatic example of how the nucleolus, a paradigmatic condensate, can exhibit complex material properties, with a radial gradient comprising both liquid-like and solid-like characteristics. Despite much speculation about the potential impact of material state on biological function, our findings provide perhaps the first clear example of condensate rheology being intimately linked to function. In particular, the constant activity of RNA Pol I transcription at the FC/DFC provides a constant source of an entangled rRNA gel, whose viscoelasticity allows it to steadily flow outward, enabling a kinetic control on the progressive processing and assembly of rRNA, prior to thermodynamic release into the nucleoplasm. This interplay between rheology, transcription, and thermodynamic driving forces is likely at play in other nuclear condensates (*50*), and may suggest novel therapeutic approaches for targeting aging, cancer, and other diseases.

## Methods

### Cell culture

HEK293, U2OS, and MV cells were cultured in 10% FBS (Atlanta biological S1150H) DMEM (GIBCO 11-965-118) supplemented with penicillin and streptomycin (Thermo Fisher 15140122) at 37°C with 5% CO_2_ in a humidified incubator. For passaging and imaging, cells were dissociated from the plate with trypsin (Trypsin-EDTA 0.05%, Fisher Scientific 25300054) and transferred to 96-well glass bottom dishes (Thomas Scientific) coated with fibronectin.

### Rodent neuron cell culture

For rodent cortical neuron culture, cortex was dissected from E17 Sprague-Dawley rat embryos (Hilltop Lab Animals Inc.), and neurons were dissociated into single cells using the Worthington Biochemical papain dissociation system. Briefly, cortices were incubated in 5 mL papain solution (20 units/mL papain, 1 mM L-cysteine, 0.5 mM EDTA, and 200 units/mL DNase in HBSS) in a 37C water bath for 20 min with no agitation. Supernatant was discarded and replaced with 3 mL inhibitor solution (1 mg/mL ovomucoid protease inhibitor, 1 mg/mL bovine serum albumin, and 167 units/mL DNase in HBSS) for 5 min at room temperature. Supernatant was discarded and replaced with another 3 mL of inhibitor solution for 5 min at room temperature. Supernatant was removed and 1.5 mL of Gibco Neurobasal Plus complete media (2% B27 Plus, 1% penstrep, 250 ng/mL Amphotericin B) was added. A flame treated pasteur pipette was used to dissociate the tissue by pipetting up and down 10 times, cell clumps were allowed to sink for 1 min, and 750 µL of dissociated cells were removed from the top and added to a new tube for subsequent steps. 750 µL more neurobasal media was added to the remaining clumped cells in the old tube, and the trituration procedure was repeated for a total of 3 dissociation steps, with all of the media being moved to the new tube after the final step. Cells were centrifuged 5 min 300g. Supernatant was discarded, and cells were resuspended in 1 mL neurobasal media for cell counting. Cells were plated with 1x CultureOne supplement (Gibco) in neurobasal media to kill glial cells. CultureOne supplement was only used in media on DIV0 (day in vitro 0), and not used in subsequent media changes. 80,000 neurons were plated per well (~40,000 cells/cm2) in 24 well glass bottom plates treated with Poly D Lysine (0.01 mg/mL overnight treated at 37C, washed x4 in PBS with no drying steps). Half of the media in each well was exchanged for fresh neurobasal media every 3-5 days. Lentivirus infection was done ~DIV13, and cells were imaged once fully mature, DIV17-21.

### iPSC cell culture

Induced pluripotent stem cells were obtained from Allen Institute for Cell Science at the Coriell Institute. The iPSC line AICS-0084-018:WTC Dual tagged FBL-mEGFP/NPM1-mTagRFPT-cl18 (mono-allelic tags) was used for our experiments. The colonies were expanded and maintained on Matrigel (Corning) in mTeSR Plus medium (Stem Cell Technologies). Cells were plated at 3000-10.000 cells per square centimeter in order to obtain ~75% confluency every 5-7 days. The cells were passaged using ReLeSR (stem cell technologies) and split at a 1:10-1:50 ratio. mTeSR plus medium was supplemented with ROCK inhibitor Y-27632 (Selleckchem) for maximum 24 hours after cryopreservation or passaging. iPSCs were cryopreserved in mTeSR Plus medium supplemented with 30% Knock Out Serum Replacement (Gibco Life Technologies) and 10% DMSO.

### Plasmid construction

DNA encoding NPM1 (Sino Biological) and RPA16 were amplified with PCR using primers synthesized by IDT. Resulting fragments were cloned into linearized FM5-mCh or FM5-GFP constructs using In-Fusion Cloning kit (Takara). The resulting plasmids were sequenced to confirm correct insertion.

### Lentiviral transduction

Using our mCh and GFP constructs, we created stably expressing cell lines transduced with lentivirus. Lentivirus was produced by transfecting the desired construct with helper plasmids PSP and VSVG into Lenti-X cells with Fugene HD transfection reagent. Virus was used to infect cell lines in 96 well plates. Three days after addition of virus, cells were passaged for stable maintenance. For rodent neurons, third generation lentivirus production was performed with standard protocols. Virus was concentrated using Lenti-X Concentrator (Takara) and resuspended in DPBS before being applied to neurons.

### Immunofluorescence

Cells were fixed by adding 4% formaldehyde to the wells. After 10 minutes, cells were washed with wash buffer (0.35% Triton-X, Thermo Fisher PRH5142, in PBS, Thermo Fisher 14190250), and permeabilized with 0.5% Triton-X in PBS for 1 hour. Cells were then blocked for 1 hour with 10% goat serum in TBS-T (20mM Tris, 150mM NaCl, 0.1% Triton-X). The primary antibody (Rabbit polyclonal anti-fibrillarin, Abcam 5821) was dissolved in blocking buffer at 0.1µg/ml and incubated overnight at 4°C. The next day, the cells were washed three times with TBS-T. The secondary antibody (AlexaFluor 647 goat-anti rabbit Thermo Fisher A-21245, 1:1000) was dissolved in blocking buffer and incubated for 2 hours at room temperature. Finally, cells were washed three times with TBS-T.

### EU labeling

For labeling transcribed RNA, the Click-iT RNA imaging kit was used (Thermo Fisher C10330). Largely, the manufacturer protocol was used, with the following adaptations. We performed the protocol in 96-well plates, with volumes adjusted accordingly. Throughout, we kept the volumes at 100µl per well. We prepared the EU solution at 2mM, and 100µl to the 100µl media already in the well, for a final concentration of 1mM. We kept the incubation of EU with the cells constant at 30 minutes. Next, we removed the media containing EU with fresh media not containing EU (for the pulse experiments). For fixation, we added 66µl of 16% formaldehyde in PBS to each well, and incubated for 15 minutes. This was followed by permeabilization with 0.5% Triton-X for 15 minutes. Addition of the Click-iT reaction cocktail per manufacturer instructions, with the exception that we use Alexa-647 (Alexa Fluor 647 Azide, Triethylammonium Salt, ThermoFisher A10277) instead of the fluorophore supplied with the kit. Reaction cocktail was incubated for 30 minutes, cells were washed once with the kit-supplied rinse buffer, and once with PBS before proceeding to imaging. For the temperature variation experiments, cells were incubated for 30 minutes with EU as described above to ensure proper incorporation. Only after EU incubation were cells moved to incubators at different temperatures.

### Actinomycin D treatment

Cells were treated for 4 hours with media containing 1 µg/mL actinomycin D (Sigma, A5156-1VL) dissolved at 0.5 mg/mL in DMSO (Sigma). Control cells were treated with media containing DMSO. EU labelling was performed as described above in media containing actinomycin D or DMSO.

### Corelet activation and FRAP

Cells expressing the two Corelet constructs (*7*) were imaged on a Nikon A1 laser scanning confocal microscope with a 100x oil immersion Apo TIRF objective (NA 1.49) and a Nikon Eclipse Ti2 body; activation was performed by imaging with the 488 laser. Following 5 minutes of activation, droplets were bleached using the 561 laser in a small region of interest and imaged at 3 seconds per frame for 5 minutes. For nucleolar frap, cells coexpressing NPM1-mCherry and RPL5-GFP were bleached using the 561nm laser for NPM1 and the 488nm laser for the RPL5 and imaged in both channels.

Images were analyzed in Fiji (ImageJ 1.52p) (*51*) and MATLAB 2019b (Mathworks). For corelets, droplets were segmented in the 488 channel; their intensities were averaged and normalized with 1 set to the frame before bleach and 0 set to the frame immediately after bleach; recovery was further normalized to non-bleached control ROIs of the same cells. For nucleoli, each nucleolus was segmented (*imbinarize*) using the non-bleached channel and the intensity profile was calculated for each nucleolus along the axis perpendicular to the half-FRAP, normalizing by the average intensity over the nucleolus. Recovery was then measured as the intensity in the half of the nucleolus that was bleached and normalized with 1 set to the frame before bleach and 0 set to the frame immediately after bleach.

### EU and immunofluorescence imaging

EU and immunofluorescence stained cells were imaged on a Nikon A1 laser scanning confocal microscope using an oil immersion objective, Plan Apo 60X/1.4NA. Imaging conditions were optimized to increase signal to noise. Proteins tagged with GFP were imaged using a 488nm laser, mCherry with 560nm, and Alexa 647 with 640nm. Images shown in Figure 2B,D, Figure 3B and Figure S3A,B were deconvolved using the Richardson-Lucy algorithm in the Nikon Elements software V4.40. All example images are optimized to show the full pixel intensity range, with the exception of Figure 2B in the EU channel.

### Sphericity analysis and Model

Cells expressing NPM1-mCherry and RPA16-GFP were imaged in three dimensions with z-stacks with a spacing of 0.3microns on a spinning-disk (Yokogawa CSU-X1) confocal microscope with a 100x oil immersion Apo TIRF objective (NA 1.49) and an Andor DU-897 EMCCD camera on a Nikon Eclipse Ti body. A 488 nm laser was used for imaging GFP and global activation, and a 561 nm laser for imaging mCherry. The imaging chamber was maintained at 37 °C and 5% CO2 (Okolab) with a 96-well plate adaptor.

Images are segmented and nucleoli are parsed as described below in the RDF calculation and analysis. Briefly, cells are manually segmented by polygon tracing and extraction, followed by automatic identification of the dense phase (“nucleolus”) as the highest intensity pixel and value after a slight blur and manual identification of a reasonable pixel and intensity to represent the dilute phase (“nucleoplasm”), and segmentation of nucleoli by utilizing these values to form a mask of the NPM1 channel. Then the identification of the true nucleolus surface and FC centers are also determined as described in the RDF analysis section below. To account for the additional blur in z for 3D data and to avoid anisotropic segmentation of the nucleolus, a slight erosion in z of 1 radial box is applied. To aid identification of the FC centers, a 2 pixel blur is applied in the x and y directions only. The nucleolar mask is then given to the function “BoundaryMesh” in Mathematica using the method DualMarchingCubes. This surface corresponds to that shown as “GC” in **Figure 2C** and **Figure S1A**.

To produce the model image, we draw from the logic of transport theory in spherical coordinates where the concentration and flux is proportional to 1/r and 1/r^2^, respectively and r is the distance from a source (which in this case is the FC center). At steady state, the flux of radiating material is balanced by a sub-saturation. To roughly incorporate these concepts into a model, we calculate the total flux at each position as a sum of 1/r^2^ over all FC centers using the same grid and spacing as the image. Then, the cut-off value for the surface (where all voxels greater than the total flux of the cut-off value are included as part of the nucleolus surface) is chosen such that the model will have the same number of voxels as the data. The boundary mesh is then produced as done with the data nucleolar mask. This surface corresponds to that shown as “Model” in **Figure 2C** and **Figure S1A**.

To approximate the degree of agreement between the two surfaces, spherical harmonics are calculated for the surface using the pseudo-inverse of the linear system with regularisation as in (*52*). Then the Pearson correlation coefficient is calculated. Note that for complicated nucleolar surfaces this analysis is not robust to reproduce the surface due to the breakdown of the 1:1 relationship between the polar and azimuth angles and the radial distance. The statistics quoted and **Figure S1B** are shown for HEK data only.

### RDF calculation and analysis

The formula for the radial distribution function (RDF) utilized throughout the manuscript is: 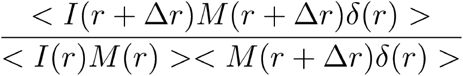; where *I*(*r*), *M*(*r*), and *δ*(*r*) are the background-subtracted image intensities (e.g. NPM1 signal), the binary mask for the region of interest (here the nucleolus), and a mask marking the pixel locations of the radial centers (here the center of the FC) at a given location r, respectively.

To calculate RDFs, confocal images of fixed cells in one z plane are taken as described above. To avoid bias, cells are first manually segmented via chosen polygon outlines blinded to the EU channel (or FIB channel for non-EU RDF experiments). Background subtraction is then performed followed by segmentation of each cell using the polygonal regions identified. The cell segmented images are then used to identify the location and intensity of the dense phase (e.g. concentration of NPM1 in the GC phase of the nucleolus) by programmatically reporting the brightest pixel location and intensity after a 2 pixel radius blur of the image to remove noise. Then looking at the NPM1 image for each cell after a 5 pixel radius blur, log transformed, and normalized to remove bias on the extent of NPM1 overexpression, a reasonable representative location of the nucleoplasm is identified and the intensity (only after the 5 pixel radius blur) is designated the dilute phase concentration for each channel. At this point, the NPM1 channel is used for each cell to form a mask by binary assignment to pixels greater than 0.25 the intensity value between the dilute and dense phase intensities followed by a filling transform, a deletion of small components with less than 300 contiguous pixels, and morphological segmentation (“MorphologicalComponents” in Mathematica) ignoring corner pixels. From these, any components less than 9 pixels large are discarded. The rest are designated as nucleoli.

For each nucleolus, the centers of FCs are identified using a combination of thresholding, distance transform, max detection, and watershedding. The nucleolus mask is then filled and eroded with a 3 pixel square to remove the expected lower intensity at the surface due to resolution limits; this mask is applied to each channel and FC centers not within this mask are discarded. Finally, to calculate the RDF, the image correlation between each channel with the image of the FC centers weighted by their relative intensity (blurred over one pixel radius for slight integration) is divided by the average intensity of the image channel. The denominator of the RDF is the image correlation between the nucleolar mask and the weighted FC centers which corresponds to the number of positions within the nucleolus at a specific distance from any FC center. To average the nucleoli, numerator and denominator RDF values at all displacements are calculated for each nucleolus and those that are smaller than the desired range of displacements are discarded; then to calculate the RDF value the the total numerator is divided by the total denominator of all remaining nucleoli. To approximate the error on the RDF value, error propagation is utilized by taking advantage of the fact that the denominator RDF corresponds to the weighted number of pixels which are being probed at a specific displacement value.

Throughout the text unless otherwise indicated the RDF points are shown with a spline fit to better depict the RDF trends and error. This is done in Mathematica by fitting a linear model with a Bernstein Basis where the input is normalized by the largest dimension in the RDF fit and raised to the 1.8 power to account for the non-linear spacing in displacement values between 0 and 1µm shown throughout the text. The number of splines utilized is 7 for the data fit between the displacements of 0 and 1µm shown primarily throughout the text. This is adjusted based on the 1.8 scaling in the displacement values and corresponds to 15 for the largest interval (2.5µm displacement) and 10 for the intermediate one (1.5µm displacement) used in the concentration dependence to account for the growing nucleolus with higher NPM1 overexpression as discussed below.

To determine concentration dependence, we fit the RDF dependence on the dense phase concentration of the NPM1 signal linearly (**Figure S5B-C)**. The shown dependence on 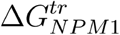 is determined as done previously (*13*) with the addition offset to account for additional background in stained cells autofluorescence that is noticeable in the trend (**Figure S5A**). At each displacement, the RDF is extrapolated to the average, minus one standard deviation, plus one standard deviation, and plus two standard deviations in the dense phase NPM1 concentration. As the 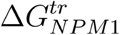 is dimensionless, we convert these dense phase concentrations into these in the text.

### Spherical diffusion-advection equation and application

To determine the flux of RNA throughout the nucleolus we utilize the incompressible advection-diffusion equation (eq 1) where D is the diffusion constant and v is the velocity.

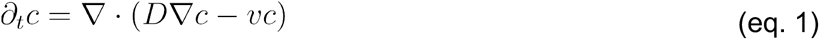

Because the radial distribution function (RDF) is spherically averaged, we can reduce the general equation to the case for spherical flow (eq. 2) dependent only on the radial distance, r, and where the flow velocity (*v*) is dependent on the injection rate (*Q*), through the relation 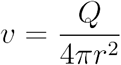. This assumes incompressibility of the RNA.

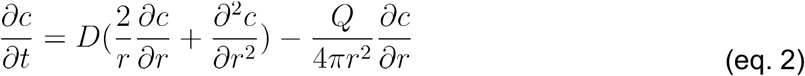

Simplifying to dimensionless units with 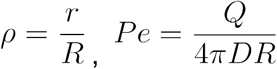, and 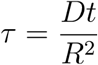, where *τ* is the dimensionless time, ρ is dimensionless radius, *Pe* is the dimensionless Peclet number describing the ratio between advection and diffusion, and *R* is the outer boundary, yields eq. 3.

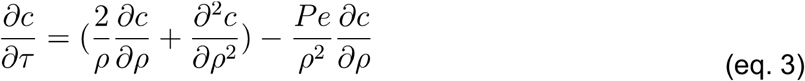

To solve eqn. 3 we use separation by parts method, also known as the fourier method (*53*). In the separation by parts method, basis solutions, *β*_*i*_(*ρ, τ*), to this equation are solved which obey the relationship *β*_*i*_(*ρ, τ*) = *χ*_*i*_(*ρ*)*ζ*_*i*_(*τ*). To determine the solutions, we start by defining the eigenvalues, *λ*_*i*_, of the basis solutions as eq. 4a which simplifies to eq. 4b.

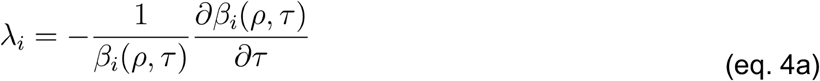

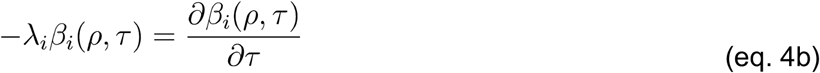

By plugging in the relationship for *β*_*i*_ and eq 3 to 4 we obtain eq. 5 and 6.

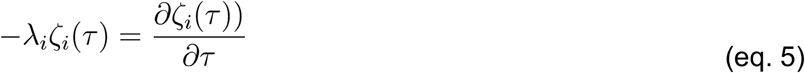

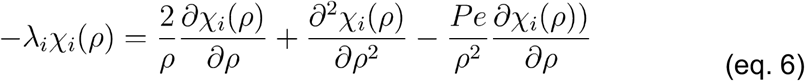

Eq. 5 can easily be solved to eq. 7.

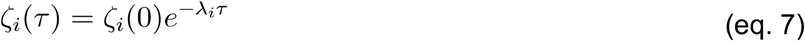

While there exists no simple form using standard commonly defined functions for the solution to eq. 6, these can be solved numerically in mathematica with the function “NDEigensolution” which yields both −*λ*_*i*_ and the numerical solution for *χ*_*i*_(*ρ*), henceforth called the eigenfunction, for the *i*th largest eigenvalue. We use a Dirichlet boundary condition equal to zero at the center and Neumann boundary condition of zero at the outer boundary (i.e. R) allowing for the constant expulsion of material due to the nature of the pulse chase experiment.

To determine the apparent Pe inside the GC phase of the nucleolus, only the smallest (i.e. index ‘zero’) eigenvalue’s eigenfunction *χ*_0_(*ρ*), or *χ*_0_(*Pe, ρ*) due to its dependence on the Peclet number, is needed; this is due to the fact that the RDF is an intensity/concentration normalized profile and thus corresponds to the apparent steady state of the RDF curve. To apply this to data, we fit the RDF for the EU (i.e. *RDE*_*EU*_(*ρ*)) data using *χ*_*i*_(*Pe, ρ*), the EU signal of NPM1 (i.e. *RDF*_*NPM*1_ (*ρ*)), and three ad-hoc parameters (*α*_*EU*_, *β, α*_*NPM*1_), that are intended to account for experimental realities such as the diffraction limit) with eq. 8.

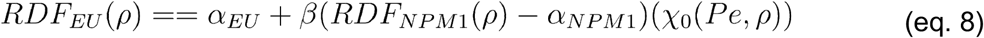

To fit the full dataset, we use R=1µm to make the displacement values into the dimensionless *ρ*. In the concentration dependent series, we adjust R by the fold increase in the maximum peak displacement in the RDF of NPM1 relative to that at no overexpression.

To determine the kinetics of the RDF profile, we solve for the 20 highest eigenfunctions and eigenvalues. Using these eigenfunctions, we solve for a linear combination of them with prefactor *ζ*_*i*_(*τ*) that yields a peaked solution near the origin at *τ*=0 and produces no significant negative value concentration profiles at all times. To convert this into an RDF we normalize the solution by the average value from the center to the outer boundary of the sphere. When showing the model in **Figure 3C**, we use unit diffusion and either a Pe of 0 or 1 for the diffusion only and diffusion plus advection cases. Furthermore in this case we use the starting conditions shown. To fit the time dependence, we fit the RDF data at *ρ*~0.9 (averaging the RDF values between the dimensionless displacements of 0.8-1) fitting for the diffusion constant D, a time offset, and baseline and scaling factors. The Pe number is set to 0.22 being the best fit to the full dataset. The outer radius, R, is set as described above.

### Half-FRAP and FRAP diffusion analysis

To determine the diffusion of nucleolar proteins following half-FRAP as described above, we will use the separation by parts as described in the previous section. Unlike the spherical transport solution, half-FRAP can be approximated as a 1D diffusion where advection can be ignored due to the fast recovery of proteins in the nucleolus. Thus we begin with the simple 1D linear diffusion equation :

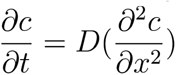

Where D is the diffusion constant. To dimensionalize the solution, we substitute 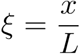 and 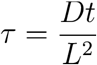 where L is the length of the nucleolus yielding:

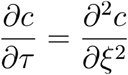

The solution to this equation is a wave (e.g. Sinusoidal). Now solving this equation from *ξ* between negative and positive half using the separation by parts yields the solution:

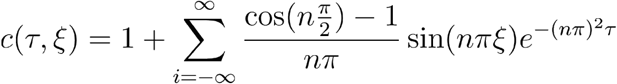

This equation is fit to half-FRAP data at *ξ*= ¼ corresponding to the location in the middle of the bleached half. Thus we fit the data using NonlinearModelFit in Mathematica to:

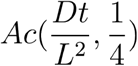

Where D, A, and L are the diffusion constant, fraction intensity recovered, and the average size of the nucleoli fit, being 4 microns. c is approximated by interpolation of the aforementioned solution with the summation truncated to between −40 and 40 using NonlinearModelFit in Mathematica.

To fit the corelet FRAP data, we fit a stretched exponential decay, 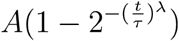, using NonlinearModelFit.

### MSD tracking

To track corelet droplets, images were taken every 3 seconds for 100 minutes following 5 minutes of initial activation on the aforementioned spinning disk as described in (*47*). For FC tracking NPM1-mcherry and RPA16-GFP were taken approximately every 20 seconds for 2.5 hours.

For all datasets, subpixel tracking was performed in TrackMate (*54*) using a Laplacian of Gaussians filter-based detector and a blob diameter of 500 nm (or an appropriate size for local activation experiments with large droplets), a threshold of 250. Trajectories were then constructed using the simple linear assignment problem (LAP) tracking with max linking and gap-closing distances of 500 nm and no frame gap accepted. Coordinates parsed into MATLAB and pair MSDs were calculated with custom routines. In mathematica, we fit the Maxwell model equation, 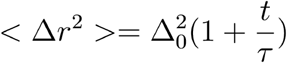, to the data.

### Fusion

To visualize corelet fusion events, large droplets were generated by local activation using a Mightex Polygon digital micromirror device (485nm) and imaged using a 561nm laser at 3 seconds per frame on the aforementioned spinning disk. To visualize nucleolar merger, images of NPM1-mCherry and RPA16-GFP were taken approximately every 20 seconds for 2.5 hours. Images of merging condensates were analyzed in Fiji and MATLAB; the aspect ratio was calculated by taking the ratio of the major and minor axes (‘*regionprops*’) at each timepoint.

## Acknowledgements

We thank the members of the Brangwynne lab, as well as Michele Tolbert, Richard Kriwacki, Lennard Wiesner, and Ned Wingreen for useful discussions. We acknowledge support from the Howard Hughes Medical Institute (HHMI), the St. Jude Collaborative on Membraneless Organelles, and grants from the Princeton Center for Complex Materials, a MRSEC (NSF DMR-2011750), and the AFOSR MURI (FA9550-20-1-0241). S.A.Q. is funded by the HHMI Hanna H. Gray Fellowship. D.S.W.L. was supported by the National Science Foundation through the Graduate Research Fellowship Program (DCE-1656466). L.A.B. is supported by an HHMI Helen Hay Whitney Fellowship.

## Author Contributions

J.A.R, J.M.E., D.S.W.L., and C.P.B. designed research. J.A.R., J.M.E., D.S.W.L., S.A.Q., L.B, L.A.B, performed research and contributed reagents. J.A.R. analyzed data. J.A.R., J.M.E., and C.P.B. wrote the manuscript with input from all authors.

## Supplementary Text

### Overlap Concentration Discussion

The concentration where polymers begin to become entangled is referred to as the overlap concentration (*27*). The overlap concentration (in units of the number of chain particles per volume) is approximated as 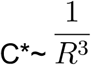 where R is the average (RMS) end-to-end distance of the chain (*27*). Given the measured end-to-end distance of the 16S (prokaryotic) rRNA (~1.5kb) of ~50 nm (*33*), the ratio of the number of basepairs or monomers between the 47S rRNA (~13.3kb) and the 16S rRNA is ~9, and the assumption that the chain acts roughly as a random walk where the end to end dimensions scale as the square root of the number of monomers, the end-to-end distance R of the full rRNA transcript is ~150 nm. Calculating the overlap concentration for this end-to-end distance yields ~300 molecules per µm^3^ or ~0.5 µM. On the other hand, full folded ribosomal subunits are anticipated to have an end-to-end distance of ~10 nm giving them an overlap concentration of ~10^6^ molecules per µm^3^ or ~1700µM. Thus the calculated overlap concentration increases over 3,000 fold during the folding of the nascent rRNA into ribosomal subunits; in other words, rRNA becomes about 3,000 fold less entangled as it goes from the nascent chain to the folded subunit.

To determine how the actual rRNA concentration compares with the overlap concentrations for both nascent and mature transcripts, we utilize curated values for the transcription rate(*k*_*trans*_~10^3.5^ transcripts/min), rRNA residence time (*τ*_*rRNA*_~45 min), average nucleolar radius (*R*_*nucl*_~1.5um), and average number of nucleoli (*N*_*nucl*_~3.5) (*55*). From the relationship 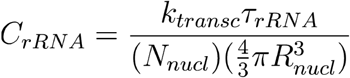, these data imply that roughly 1.5*10^5^ rRNA molecules dwell within the total average nucleolar volume of 50 µm^3^, yielding a concentration ~3000 molecules per µm^3 or ~5µM. Since this is higher than the overlap concentration for nascent rRNA chains, i.e. 5µM>0.5µM, this implies that they are significantly entangled, as can be visualized in Figure 2A. On the other hand, since the rRNA concentration is much less than the overlap concentration for folded rRNA subunits, i.e. and 5µM<<2,000µM, assembled ribosomal subunits are certainly not entangled. The nascent 13.3kb rRNA transcript is cleaved into several fragments, including the 5.8S (length=160bp), 28S (length=4.7kb), and 18S (length=1.9kb), which can partially help disentangle the chains. However, since the overlap concentration scales with the inverse physical size of the chain R (i.e. R_ee_^3^ or R_g_^3^), and making the simplifying approximation that the chains behave as random coils yields a dependence of R~N^2^, there is a relatively weak dependence of the overlap concentration on the number of nucleotides comprising the chain, i.e. C*~N^-⅔^. As a consequence, folding of the chain, i.e. going from a random-coil to a more compact structure dictated by its intra- and inter-molecular interactions, is likely the more important contribution to the progressive loss of rRNA entanglement.

### Supplementary Figures

**Figure S1:**
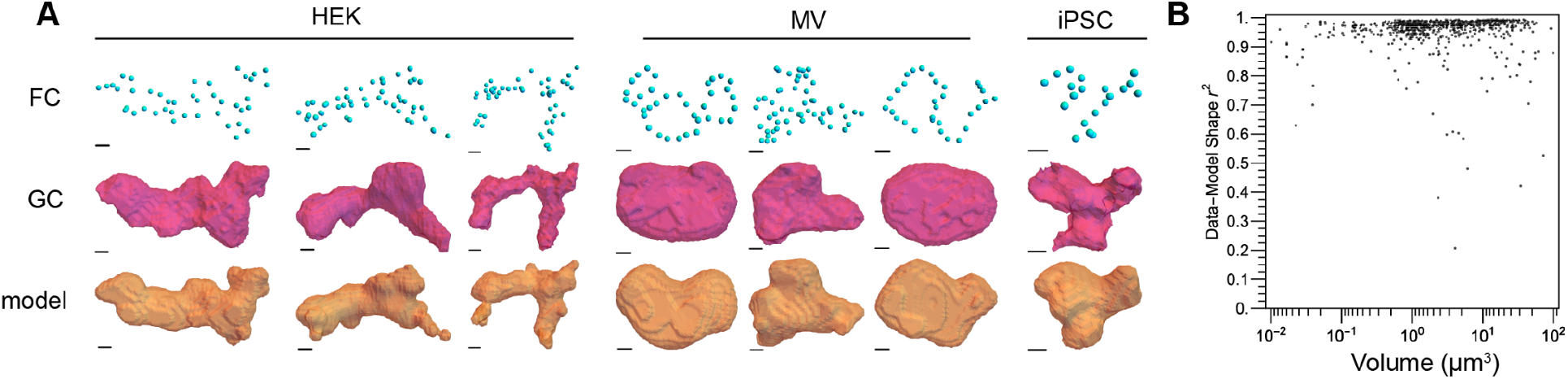
Additional examples for nucleolar shape analysis. A) NOR location is sufficient to describe nucleolar shape, additional examples in multiple cell types from **Figure 2C**. B) Spherical harmonic decomposition shows strong agreement (R^2^ = 0.96 ± 0.01) between data (i.e., GC segmented surface) and model (i.e., FC derived nucleolar shape) with little dependence on the total volume of the nucleoli.

**Figure S2:**
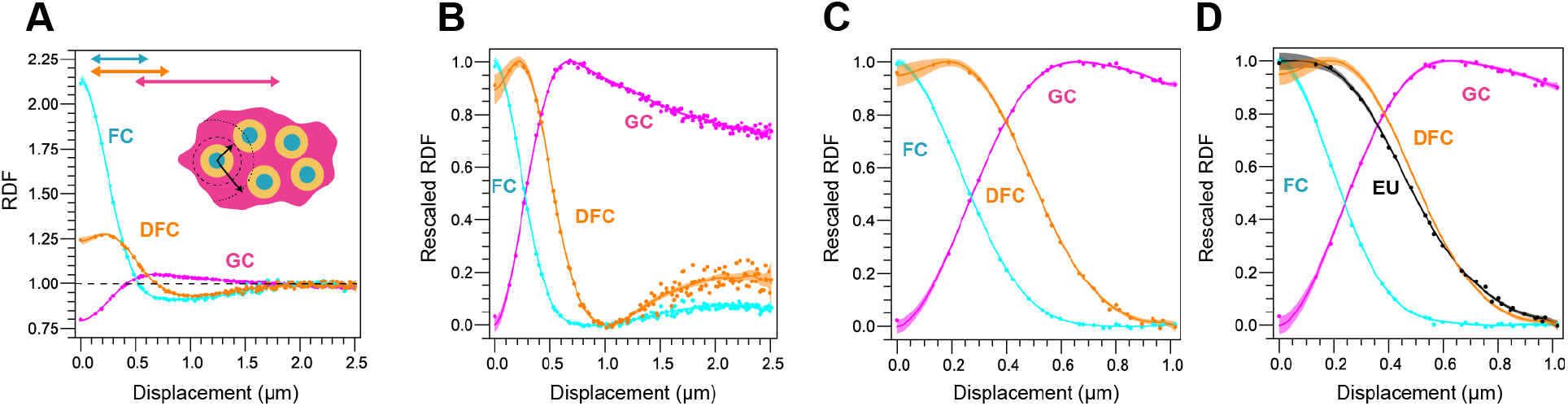
Stepwise RDF calculation. A) Same as Figure 2E, the RDF of nucleolar phases follows concentrically. B) Rescaled RDF, with the DFC curve exhibiting a minimum point at ~1µm, indicating where adjacent FCs start to contribute to the RDF. C) RDF focused on the relevant part <1µm (here, N=333 nucleoli; this is increased due to the inclusion of smaller nucleoli from the smaller displacement maximum, see methods). Unlike **Figure 2F**, here all data is from the same experiment. D. Same as **Figure 2F**, for comparison to **Figure S2C**.

**Figure S3:**
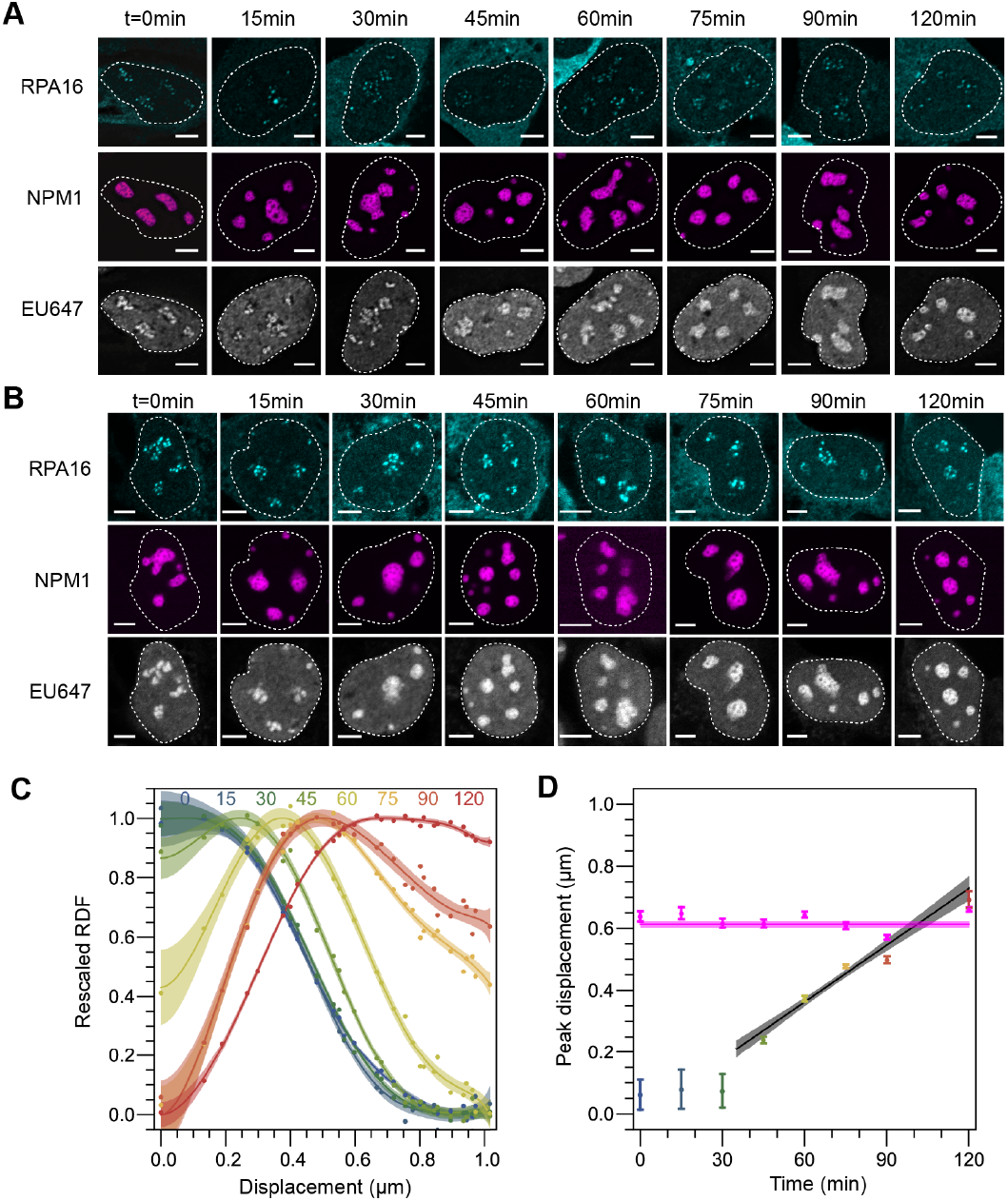
Replicates of pulse chase experiment in A) Nuclei showing EU dynamics over time. Nucleoli in Figure 3B are taken from this series. B) Replicate of experiment shown in A. C) Rescaled RDF quantification of EU signal from replicate shown in B (N= 60, 55, 52, 55, 92, 61, 66, and 77 nucleoli for sequential timepoints). D) RNA peak over time where linear fit is ~1Å/s as with **Figure 3E**

**Figure S4:**
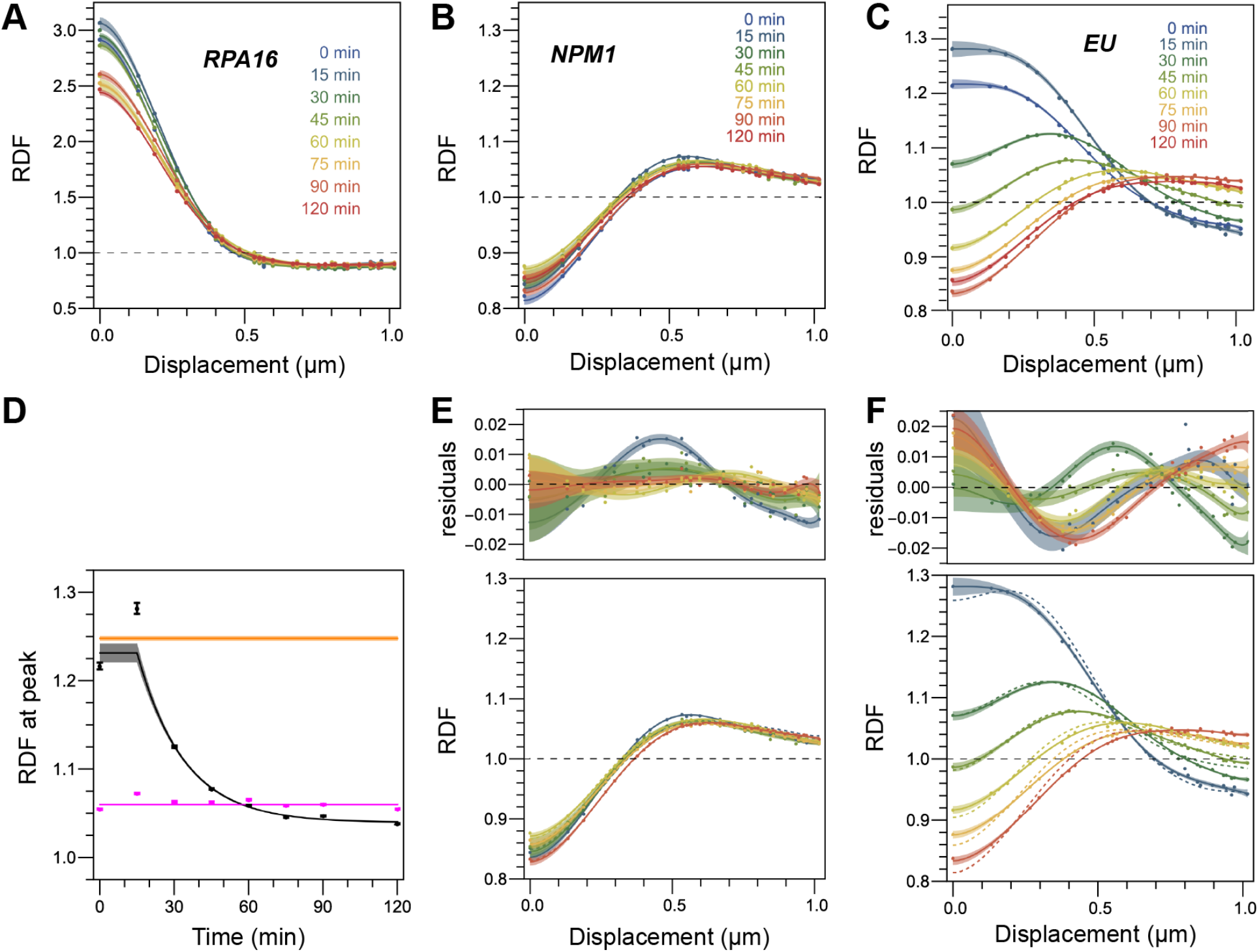
RDF validation. A) The RDF of RPA16 over time. B) The RDF of NPM1 over time. C) The RDF of EU over time (same as **Figure 3C**). D. The RDF at the peak is stable for NPM1 (magenta), but decreases for EU (black). Shown in orange is the RDF value at the peak for the DFC. E-F) Rapid-partitioning model between the DFC and GC (i.e. linear combination of basis RDFs for DFC and GC) reasonably fits the NPM1 data (E) but fails to fit the EU data (F). Top, residuals between data and model with error curves and fit lines using spline fit to data. Bottom, dashed lines show best fit from model compared to data and spline fit to data.

**Figure S5:**
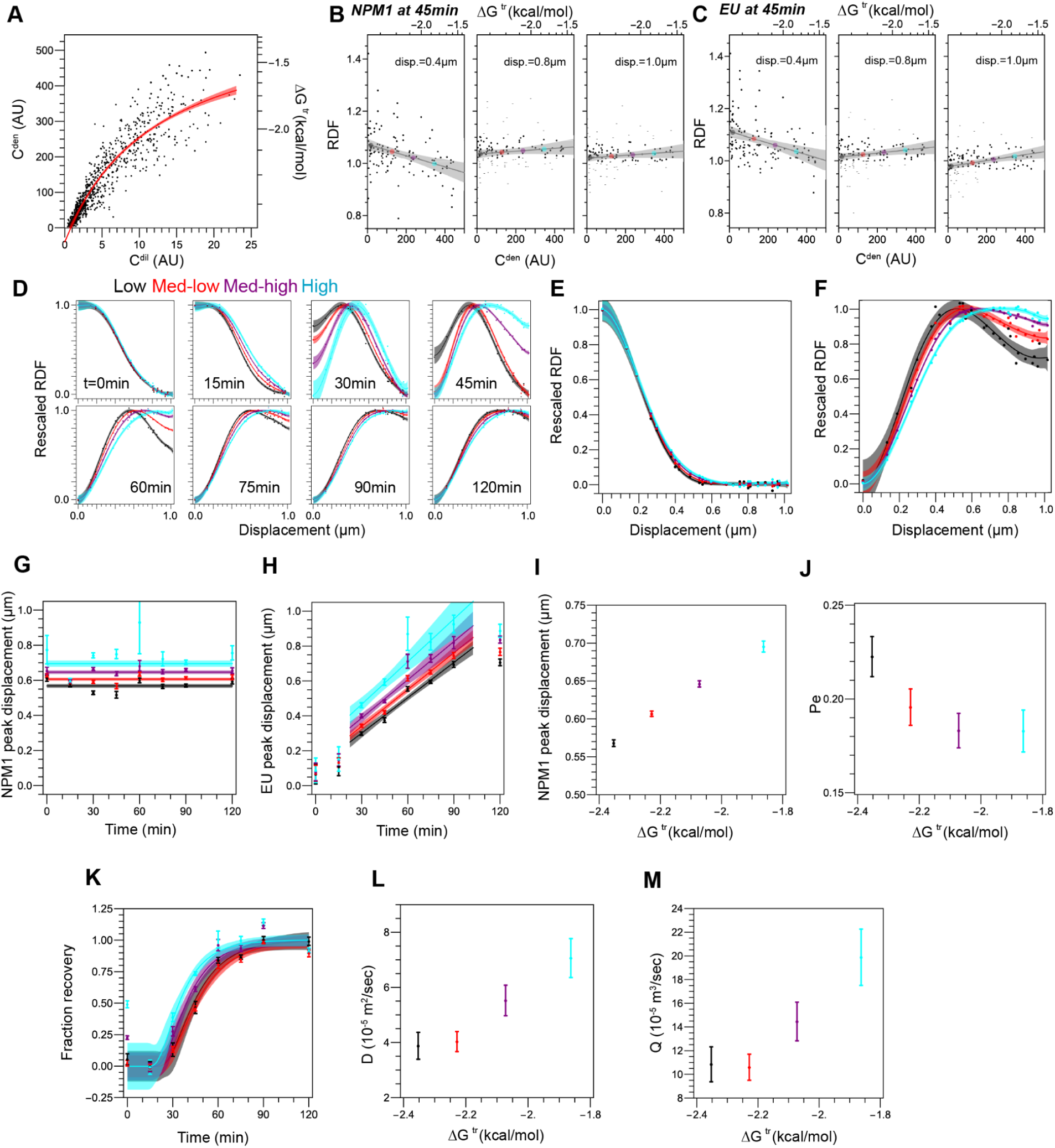
NPM1 concentration dependence on RDF curves. A) Dependendence of NPM1 dense and dilute concentrations with NPM1 overexpression. Corresponding approximate ΔC^tr^ shown on the right axis. B) RDF of NPM1 at t=45min at different displacements for different NPM1 concentrations. C) RDF of EU at t=45min at different displacements for different NPM concentrations. For B and C, dots correspond to concentration of NPM1 used for subsequent labels of the degree of NPM1 overexpression. D) Rescaled RDF of EU at different timepoints for different NPM1 concentrations. E-F) NPM1 concentration dependence of rescaled RPA16 (E) and NPM1 (F) RDF at 45min; these show the lack and significant dependence of FC and GC structure, respectively on NPM1 concentration. G) NPM1 peak displacement over time at different NPM1 concentrations. H) EU peak displacement over time at different NPM1 concentrations. I) NPM1 peak displacement from (G). J) Peclet number dependence on NPM1 concentrations. K) NPM1 concentration dependence on the EU movement to the edge of the GC. L-M) Diffusion constant (L) and advection (M) dependence on NPM1 concentration.

**Figure S6:**
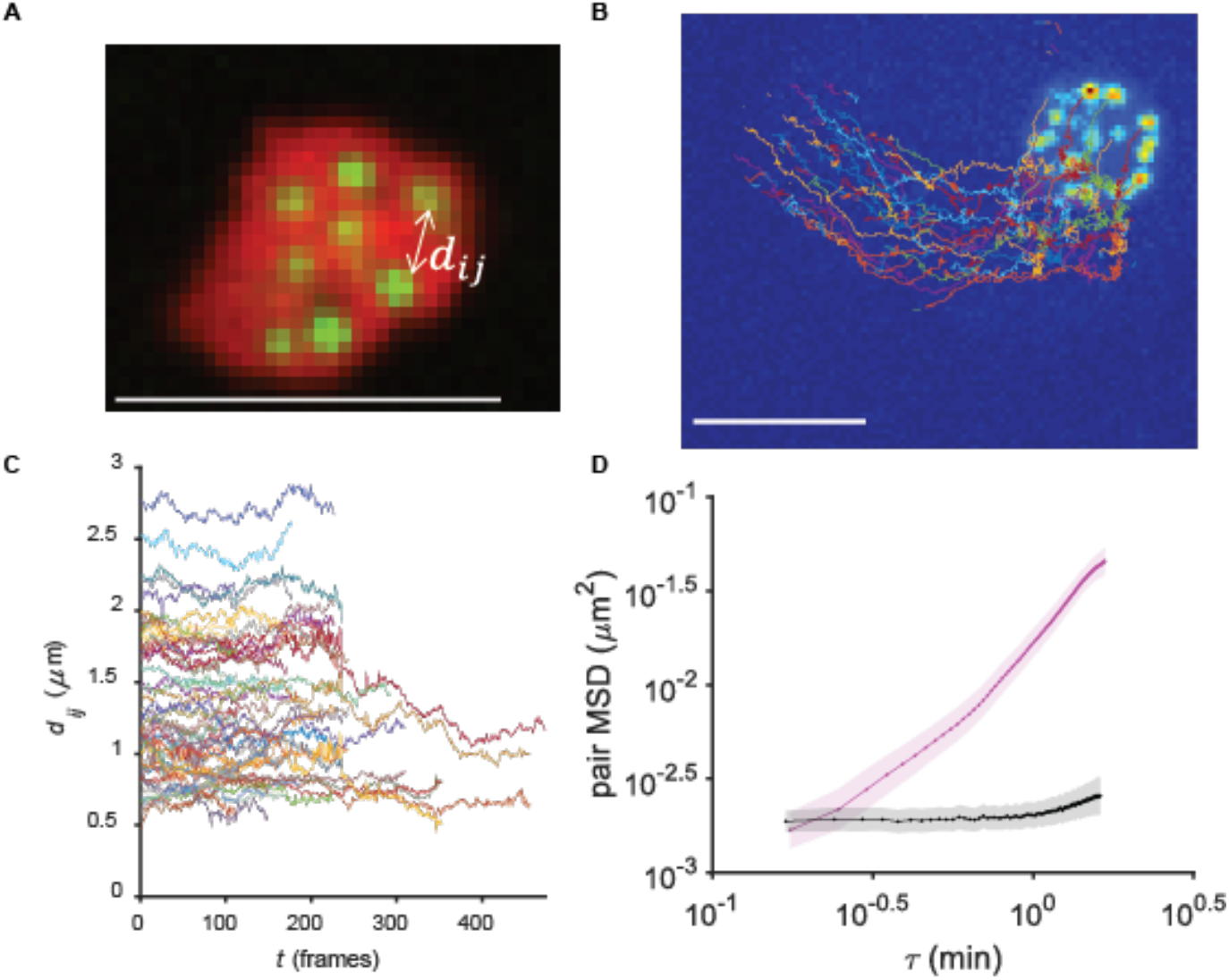
pair-MSD calculation details and controls. **A)** To characterize NOR dynamics, cells co-expressing RPA-GFP and NPM1-mCherry were imaged for 2.5 hours at an interval of 20 seconds per frame and individual nucleoli were analyzed as shown. Scale bar is 5µm. B) Single particle tracking of RPA foci revealed bulk motion associated with individual nucleoli and nuclei. C) In order to correct for this, following (*44*), the displacement was computed for each individual displacements *d*_*ij*_ (as shown in **Figure. S6A**) for each possible pair of foci. D) The pair MSD was calculated from pair displacements *d*_*ij*_ and averaged over all pairs within individual nucleoli and then over the population of nucleoli. Pair trajectories with fewer than 100 continuous frames and time lags of fewer than 90 frames were excluded from the analysis. Noise floor calculating for pair MSD was calculated by tracking pairs of RPA foci in fixed U2OS cells (531 pairs in 20 nucleoli), plotted in black. HEK pair MSD (508 pairs in 7 nucleoli) presented in main text plotted in magenta, demonstrating that the first two points are near noise floor but remaining lags are far above it, suggesting that NORs dynamics transition from a subdiffusive to a diffusive regime.

